# Ongoing convergent evolution of a selfing syndrome threatens plant-pollinator interactions

**DOI:** 10.1101/2023.05.25.542270

**Authors:** Samson Acoca-Pidolle, Perrine Gauthier, Louis Devresse, Antoine Deverge Merdrignac, Virginie Pons, Pierre-Olivier Cheptou

## Abstract

- Plant-pollinator interactions evolved early in the angiosperm radiation. Ongoing environmental changes are however leading to pollinator declines that may cause pollen limitation to plants and change the evolutionary pressures shaping plant mating systems.
- We used resurrection ecology methodology to contrast ancestors and contemporary descendants in four natural populations of the field pansy (*Viola arvensis*) in the Paris region (France), a depauperate pollinator environment. We combine population genetics analysis, phenotypic measurements and behavioural tests on a common garden experiment.
- Population genetics analysis reveals 27% increase in realized selfing rates in the field during this period. We documented trait evolution towards smaller and less conspicuous corollas, reduced nectar production and reduced attractiveness to bumblebees, with these trait shifts convergent across the four studied populations.
- We demonstrate the rapid evolution of a selfing syndrome in the four studied plant populations, associated with a weakening of the interactions with pollinators over the last three decades. This study demonstrates that plant mating systems can evolve rapidly in natural populations in the face of ongoing environmental changes. The rapid evolution towards a selfing syndrome may in turn further accelerate pollinator declines, in an eco-evolutionary feedback loop with broader implications to natural ecosystems.

## Introduction

The astonishing diversification of angiosperms, which started 100 million years ago (Benton *et al*., 2022), is commonly thought to have been stimulated and hastened by the diversity of plant-pollinator interactions (Whittall & Hodges, 2007). More than 80% of extant angiosperms rely on animals for pollination (Ollerton *et al*., 2011), thus plants and pollinators have a close relationship. Yet, previous studies have shown that a decline of pollinators (including bees, Anthophila) is ongoing across the world (Goulson *et al*., 2015; Hallmann *et al*., 2017; Grab *et al*., 2019). A lack of pollinators can in turn directly affect plant reproduction; indeed parallel declines in insect-pollinated plants and insects have been documented, over the last 50 years (Biesmeijer *et al*., 2006). At evolutionary time scales, pollen limitation has been shown to represent a major selective pressure in the evolution of self-fertilization (selfing) in theoretical evolutionary models (Lloyd, 1992; Thomann *et al*., 2013). In recent years, such a shift towards selfing has been experimentally reproduced in the absence of pollinators (Bodbyl Roels & Kelly, 2011; Gervasi & Schiestl, 2017; Ramos & Schiestl, 2019) and illustrated in natural populations impoverished in pollinators (Brys & Jacquemyn, 2012). This evolution of selfing is commonly associated with changes in reproductive traits such as a decrease in the distance between male and female organs (herkogamy) (Bodbyl Roels & Kelly, 2011; Brys & Jacquemyn, 2012; Ramos & Schiestl, 2019) but also a decrease in floral traits mediating plant-pollinator interactions, such as a decrease in corolla and petal size, and decreased scent and nectar production (Brys & Jacquemyn, 2012; Gervasi & Schiestl, 2017; Ramos & Schiestl, 2019). A shift in mating system and traits leading to higher selfing, less conspicuous and attractive flowers has been well described at the species level (Stebbins, 1957; Sicard & Lenhard, 2011). This set of trait changes characterizes the evolution of a selfing syndrome (Ornduff, 1969; Sicard & Lenhard, 2011).

Faced with pollinator declines, natural populations are expected to evolve towards reduced plant-pollinator interactions and increased autonomous selfing (Thomann *et al*., 2013). However, demonstrating trait evolution requires distinguishing between the genetic and the environmental components of phenotypes, which is not straightforward in longitudinal studies of natural populations. The “resurrection ecology” approach, which involves growing dormant seeds collected from field populations multiple generations ago alongside their naturally occurring descendants (Franks *et al*., 2007; Weider *et al*., 2018), is a powerful methodology to detect such evolution (Weider *et al*., 2018). Indeed, this approach enables the contrast between traits of ancestral versus recent genotypes in a common environment, thus revealing the genetic components of temporal trait variation. Here, we took advantage of an ancestral seed collection of the field pansy (*Viola arvensis*) collected about two decades ago in the Paris region (France) to test whether this species’ mating system has evolved in response to recent pollinator declines.

The study species is an annual weed of intensive crops (mostly wheat and oilseed rape) that produces only chasmogamous, zygomorphic, showy flowers and mixed selfing rates (Scoppola & Lattanzi, 2012; Cheptou *et al*., 2022). The length of the nectar spur and observations *in natura* are indicative of pollination by long-tongued bees. For example, *Bombus terrestris* is one of the wild pollinators observed. The study region has degraded pollination services compared with the rest of France (Martin *et al*., 2019). The status of pollinators or pollination services in such intensive agricultural areas has been reported as declining in several studies over our timescale (Goulson *et al*., 2015; Grab *et al*., 2019; Potts *et al*., 2010; Burkle *et al*., 2013; Raven & Wagner, 2021; Janousek *et al*., 2023). Parts of Belgium (some 100 to 200 km north of our study area) have experienced similar agricultural dynamics as in our study region, and studies there indicate that 32.8% of bee species are threatened or extinct based on comparisons with their presence before and after 1970 (Drossart *et al*., 2019). These studies also highlighted a more severe decline or higher vulnerability of long-tongued bees (Biesmeijer *et al*., 2006; Goulson *et al*., 2015; Drossart *et al*., 2019; Janousek *et al*., 2023).

In this study, we contrast ancestors (collected in the 1990s to early 2000s) and contemporary descendants (collected in 2021) from four localities across a range of 100 kilometres in the study region, combining genetic and phenotypic approaches. After a refresher generation (F0), phenotypic traits were measured in the second generation (F1) in 2022 in an air-conditioned greenhouse. We quantified the evolution of traits mediating plant-pollinator interactions as well as other morphological traits not directly involved in plant-pollinator interactions. We also performed a pollinator preference experiment by monitoring bumblebee visitations in mixed artificial populations containing genotypes of the ancestral and descendant populations. Together, these approaches allowed us to document on the evolution of plant-pollinator interactions and evolution of plant mating system induced in recent environmental changes.

## Material and Methods

### Study system and sampling

*Viola arvensis* Murray is an annual species of wild pansy (Violaceae). It is commonly found in fields as a weed or in meadows. This species is described as self-compatible with a mixed mating system (Scoppola & Lattanzi, 2012; Cheptou *et al*., 2022). Plants were sampled twice in agricultural fields in four locations of the Parisian Basin (see Table **S1**). The first sampling was done and conserved by the Conservatoire Botanique National de Bailleul for Crouy (Cr) and Lhuys (Lh) in the 1990’s and by the Conservatoire Botanique National du Bassin Parisien for Commeny (Co) and Guernes (Gu) in the 2000’s. Seeds were collected by sampling fruits on a minimum of 100 mothers picked randomly in each population. The second sampling was performed in exactly the same locations, in an area that overlapped the ancestral sampling area as much as possible, in February 2021 by randomly sampling 40 or more seedlings. The minimal sampling areas was of 2,000 m^2^ and correspond to cases were all the population is circumscribed on this area. This sampling scheme produced four couples, one for each location, of ancestral (A) and descendant (D) populations, thus height populations in total.

### Experimental design

To avoid maternal and potential storage effects (Franks *et al*., 2018), a refresher generation (F0) was grown in common garden with plants (A and D) originated from the field (Fig. **S1**). As ancestral populations were stored seeds and descendant ones were collected at seedling stage in the field, common conditions for the F0 started at seedling stage. For ancestral populations, in mid-November 2020, seed lots were placed in germination chambers in petri dishes with vermiculite under light at 15°C for 12h and in the dark at 6°C for 12h. Germination rates were of 50% to 65%. 32 seedlings of ancestral and descendant populations were transplanted in late February 2021 in the field of the Baillarguet site of the experimental platform “Terrains d’Expériences” of the Labex Cemeb, 10 km north of Montpellier, France (43°40′53.92′′N, 3°52′28.74′′E). A single event of mortality was observed in CrD population. Populations were separated in insect-proof mesh cages (Fig. **S1**). During full blooming, in mid-April, we introduced commercial bumblebee hives, Natupol from Koppert France® (Cavaillon, France), into the cages to produce open-pollinated fruits that we collected.

These collected fruits produced by the refresher generation, in bumblebee pollination in common environment, were used for the experimental generation (F1). We randomly selected 20 mothers per population in the F0 generation to construct 20 families in the F1 generation. In December 2021, 30 seeds per family were placed in germination chambers in petri dishes with vermiculite under light at 15°C for 12h and in the dark at 6°C for 12h, to obtain enough plants for measurements. Germination rates per population range between 75% and 90%. In each family we selected five seedlings to establish families, each composed of five siblings (Fig. **S1**). Three families did not germinate enough to obtain five plants and thus were composed of one, two or four siblings. Due to the high selfing ability of *Viola arvensis*, most of the individuals were inbred, for both ancestors and descendants (confirmed by F1-individuals microsatellite genotyping, unpublished results). Plants were placed in individual pots (11 × 11 × 11 cm^3^) filled with standardized soil in late February and randomized in a greenhouse of the experimental platform “Terrains d’Expériences” of the Labex Cemeb, Montpellier, France. Watering was manual and done when needed. In total, 792 plants were followed during this experiment and no early mortality was observed. When all the morphological measurement were performed, plants were moved outside to complete their life cycle.

### Population genetics

32 F0-plants per population were genotyped using eleven microsatellite markers, 6 designed for *Viola arvensis* (Cheptou *et al*., 2022) and 5 designed on *Viola tricolor* (Latron *et al*., 2018), a closely related species (Scoppola & Lattanzi, 2012). DNA was extracted from fresh leaves using Dneasy Plant Mini Kit from QIAGEN®. Analyses of metrics such as allelic richness, expected heterozygosity (*H_e_*), using the R package genepop (Rousset, 2023), were performed to look at a change in genetic diversity over time. A drastic decrease in allelic richness (or *H_e_*) is indicative of genetic drift, strong population bottlenecks or a sampling bias. For the study species, a widespread weed, reduced population sizes can be ruled out because weed surveys have revealed higher occurrence in agrosystems in the study region (Fried & Reboud, 2007). Heterozygosity deficiency (*F*_IS_) analyses were performed, using the R package HierFstats (Goudet *et al*., 2022), on eleven microsatellite markers. *F*_IS_ was used to estimate the selfing rate using the following formula: 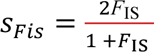 (Crow & Kimura, 1970). This estimate of selfing integrates inbreeding over several generations which buffer annual effects. Confidence intervals were obtained by bootstraps of 32 individuals per population and calculation of *F*_IS_ (50,000 per population). Bootstraps allowed us to record differences between descendant and ancestral *F*_IS_ for each locality. The number of cases where the difference in reconstructed *F*_IS_ is inferior or equal to 0 divided by the number of bootstrap (50,000) gives us a *P*-value for H_0_: “True difference < 0 or True difference = 0”. We performed a Fisher’s combined probability test on the *P*-values obtained for each locality (Fisher 1970) to test for a global increase in selfing rate between ancestral and descendant populations. Because some *P*-values obtained were equal to 0, we had 1/50,000, the minimal *P*-value measurable by this method, to each *P*-value to compute the Fisher’s combined probability test (conservative test).

### Measurements of plant traits

Plant traits were measured on the test generation (F1). All the morphometric measurements (Fig. **S2**) were performed by a single experimenter during all the experimentation, to avoid any experimenter effect, using a digital caliper to 0.01 mm.

Corolla length, labellum width, spur length, sepal length, number of guides and anthesis duration were measured on the five first opened flowers of the main stem of each plant. We chose the first five opened flowers of the main stem because more exhaustive measurements in the F0 showed that they capture the floral variance of an individual. Measurements were made day+2 after anthesis, or day+3 when falling on weekends. The opening of flowers was followed every working day to assess the anthesis duration. Floral area approximation is the product of corolla length and labellum width. Sepal/corolla length ratio is the ratio of sepal to corolla lengths. Around 4,000 flowers were measured. Flowering date is the date of anthesis of the first flower of a plant. The number of flowers opened per plant was counted each working day, from the opening of the first flower, for one month and half, and two more times with one week and then two weeks delay between counts. This number was integrated using trapezoidal rule and divided by the duration between the flowering date of the plant and the last day of measurement, giving the average floral display. Nectar volume was collected and cumulated from three flowers per plant using 0.5 μL microcapillaries. It was measured randomly on individuals of the two populations of the same location simultaneously to avoid differences in nectar production caused by the time of measurement. Thus, for this measure, locality effect (see “Statistical analysis”) is confounded with the day of measurement. It was measured on half of the plants from each population, representing all the families, thus 400 plants were sampled.

Vegetative traits such as rosette diameter, leaf length and leaf width were measured at the flowering date of the plant. Leaf length and leaf width were measured on the biggest leaf at this time, and the product of these two measures gives leaf area approximation. Biomass was measured on dried naturally dead plants at the end of their life cycle.

To measure production of seeds in absence and in presence of pollinators, when floral and leaf traits had been scored, youngest receptive flowers and oldest non-receptive ones on the same stem were marked before the exposition to pollinator to differentiate fruits produced in zero-pollinator context (self-pollination) versus fruits produced in open pollination. Thus, open pollination fruits were produced later in the stem. Then, plants were placed by groups of twenty in insect-proof mesh cages which contained commercial *Bombus terrestris* bumblebee hives, Natupol from Koppert France® (see “Bumblebee preference experiment” below). After 2 hours minimum (including the time for the bumblebee preference experiment), plants were moved outside in open pollination context. For each plant, just before the dehiscence of the marked fruit produced by self-pollination, we sampled the two fruits marked before (produced without or with exposition to pollinators) at the same time and recorded the difference in ranks of the fruits on the stem (minimum 1, by definition, to maximum 5, with a mean of 1.6). The sampling of the two fruits was performed at the same time because of the practical difficulty to follow 1,600 fruits independently. Thus, open pollination fruits are not fully developed and comparing weights between open pollination and self-pollination fruits is slightly biased. However, comparing each measure between populations is fully possible while differences in floral positions are distributed randomly between populations. Reproductive traits were obtained while counting and weighing all the seeds contained by fruit to the nearest 0.1 mg using a precision balance. The seed weight reported is the measure of the weight of seeds contained in the fruit divided by the number of seeds counted in the fruit. The ratio of the number of seeds produced by self-pollination to the number of seeds produced by open pollination is a proxy of pollen limitation due to the absence of pollinators, the less this limitation is the more the self-pollination ability is.

### Bumblebee preference experiment

Once all the floral trait measures have been made, F1-plants were exposed to commercial bumblebee (*Bombus terrestris*) hives, Natupol from Koppert France®, in two insect-proof mesh cages. In each insect-proof cage, we placed, for the entire experiment, a bumblebee hive of around 30 workers with a queen, with nutrient reserves. We randomly arranged plants from the same location in a group of 20 plants, composed of 10 plants from each ancestral and descendant population randomly selected in one of the two insect-proof cages. Pots were equally spaced at 35 cm. The observer, stationed in the entrance of the insect-proof cages, selected a foraging worker bumblebee, and recorded each flower visit during 10 to 15 minutes. Cage number (1, 2), position in the cage (1-20) and plant identities were also recorded. Experiments were performed between the 2^nd^ and the 19^th^ of May 2022. We performed the experiment on 8 groups for each locality but just 6 and 7 observations were kept for Lhuys and Crouy respectively, because worker bumblebees foraged less than 10 minutes in the discarded observations. Of the 580 plants in our dataset, 3 are individuals exposed two times to reach the 20 individuals per observation. During a 10-15 minutes test, 4.28 plants were not visited on average and 124 in total for all the tests. In the analyses, we sorted out plants visited and unvisited (see below). We calculated the proportion of visits per plant as the ratio between the number of visits on a given plant divided by the total number of visits during a flight.

### Statistical analysis

Changes in traits were evaluated using linear discriminant analysis (LDA) and univariate linear mixed models (LMMs). We performed the LDA over nine traits supposed to be linked with reproductive system; corolla length, labellum width, floral area, number of guides, anthesis duration, flowering date, volume of nectar, average floral display and ratio of seeds produced in self-pollination to open pollination. For traits with repeated measures, we used the means of traits per individual. We normalized values by locality in order to focus on age effect, *i.e.* ancestral or descendant. We predefined only the population (8 groups) in the LDA and not the age. LDA gives linear combinations of traits (discriminant functions) that attempt to capture differences among groups. In our case, it produced seven discriminant functions. The two first functions represent 55% and 22% of the variation.

Univariate analyses were conducted using LMMs with each trait as response variable, age (ancestral or descendant), locality (4 localities), and the interaction between age and locality as fixed factors, and family as random factor nested within the combination of age and locality. In order to account for repeated measures on the same individual for floral measurement, we add individual as random factor nested in family. For seeds traits, we accounted for the difference in flower position between the two fruits collected adding this difference as random factor nested in family. When there was significant age effect, analysis was followed by T-test comparison, on the means of the trait per individual for repeated measures, of ancestral and descendant populations of each locality. T-test *p*-values were corrected using Bonferroni correction. Rosette diameter, leaf length, width and area, nectar volume and ratio of seeds without/with pollinators were log-transformed to approach normal distribution of residuals. In these analyses, age significant effect represents a signal of global evolution of traits between ancestral and descendant populations, locality significant effect, represents a difference in traits between localities. A significant effect of the interaction between age and locality represents a difference in the magnitude or in the sign of the evolution between localities. In the case of a significant age effect and a significant interaction effect, if evolution is consistent (same sign of evolution) among localities and only the magnitude change, we can hypothesize a directional selective force that selects a trait in a similar direction in every locality. In other cases, a significant interaction between age and locality indicates local differences in selective pressures on the trait, in genetic variance of the trait or alternatively genetic drift.

For bumblebee preference analyses, we first performed a GLMM to test the effect of age (ancestral or descendant) on the number of unvisited plants (unvisited plant proportions: CoA: 0.24, CoD: 0.35; CrA: 0.26, CrD: 0.26; GuA: 0.15, GuD: 0.18; LhA: 0.13, LhD: 0.12). We modelled it with age and locality as fixed effect, family, cage, position nested in cage and date as random effects, with binomial distribution and logit link, as response variable was a Boolean variable with TRUE for visited plants and FALSE otherwise. There was no significant effect of age in the probability of visit (N = 580 plants, estimates: Ancestral-Descendant = 0.223, *P* = 0.296). Thus, to avoid non-normal distribution, we excluded unvisited plants (N = 124) in the rest of the analysis (similar results were found when including or not unvisited plants). We performed a LMM with log-proportion of visits per plant as response variable (N = 456), and the same structure as for other traits with age, locality, and the interaction between age and locality as fixed effect, family as random effect and we added cage, position nested in cage and date as random effects. We then performed T-test comparisons between ancestors and descendants for given locality and corrected *p*-values using Bonferroni correction.

All our statistical analyses were performed using the R software (R Core Team, Version 4.2.1). LDA was performed using the “lda” function implemented in the package “MASS” (Venables & Ripley, 2002). LMMs and GLMM were performed using the “lmer” and “glmer” functions implemented in the package “lme4” (Bates *et al*., 2015).

## Results

The localities exhibited no overall significant differences in allelic richness, except a slight decrease in Commeny, nor genetic diversity between ancestral and descendant populations based on eleven microsatellite markers (Table **1**). In the case of a resampling bias of the populations or a strong demographic bottleneck or genetic drift between the two samples, we predicted a decrease in allelic richness and genetic diversity. Our results do not support these scenarios. In contrast, we did observe a consistent increase in selfing rates in all localities between ancestral and descendant populations, except for Crouy (Table **1**), with an average and significant increase of 27% (Fisher test, 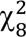 = 45.81, *P* < 0.001).

**Table 1.** Allelic richness, genetic diversity and selfing rates in our populations (mean ± SD or [95% CI]) of *Viola arvensis*. Average allelic richness and genetic diversity are based on eleven microsatellite markers. These markers were genotyped on the thirty-two individuals per population of the F0 generation issued from natural populations. For selfing rate, confidence intervals (CI) were obtained by bootstraps on individuals (50,000). Fisher’s combined probability test gave a significant overall increase in *F*_IS_ (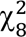 = 45.81, *P* < 0.001, see Material and methods). The first two letters are the name of the locality (Co = Commeny; Cr = Crouy; Gu = Guernes; Lh = Lhuys). “A” ancestral population (collected in 2000 for Co, 1993 for Cr, 2001 for Gu and 1992 for Lh) and “D” descendant population (all collected in 2021).

| Population | CoA | CrA | GuA | LhA |
| --- | --- | --- | --- | --- |
|  | CoD | CrD | GuD | LhD |
| Average allelic richness | 2.55 ± 1.21 | 3.09 ± 1.81 | 2.64 ± 1.29 | 2.82 ± 1.78 |
| | 1.73 ± 0.79 | 3.36 ± 1.86 | 3.27 ± 1.56 | 2.18 ± 1.08 |
| Genetic diversity ($H_e$) | 0.302 ± 0.272 | 0.436 ± 0.264 | 0.314 ± 0.210 | 0.315 ± 0.291 |
| | 0.099 ± 0.133 | 0.432 ± 0.252 | 0.324 ± 0.255 | 0.245 ± 0.225 |
| Selfing rate [95% CI]<br>derived from heterozygosity<br>deficiency, $F_{IS}$ | 0.74 [0.57-0.85] | 0.66 [0.46-0.79] | 0.44 [0.22-0.64] | 0.33 [0.09-0.48] |
|  | 0.89 [0.66-1] | 0.70 [0.55-0.80] | 0.80 [0.71-0.88] | 0.88 [0.78-0.95] |

Descriptive multivariate analysis performed on eight floral traits associated with the mating system (corolla length, labellum width, floral area, number of guides, anthesis duration, flowering date, volume of nectar and average floral display) and the ability of plant to set seeds in the absence of pollinators (*i.e.* autonomous self-pollination), corrected for differences between localities, show differentiation between ancestral and descendant populations. We chose these traits because they are usually associated with shifts in mating systems (Sicard & Lenhard, 2011; Bodbyl Roels & Kelly, 2011; Gervasi & Schiestl, 2017; Ramos & Schiestl, 2019). The descendant populations and ancestral populations form two distinct clusters in the trait space (Fig. **1**), revealing convergent directional changes of floral traits in all four localities. Linear models on the nineteen traits measured revealed consistent patterns (Table **2**). We detected a significant difference between ancestral and descendant populations for all the floral traits except sepal length (Table **2**). Plants have evolved a 10% decrease in floral area (Fig. **2a**) through a combined decrease in the labellum width and flower length (Table **2**). Flowers also have fewer nectar-guides (Fig. **2b**). In contrast, none of the morphological traits presumed to be unlinked to pollinator attraction (vegetative traits and sepal length) showed consistent changes (Fig. **2c**, Table **2**). The increase in sepal/corolla ratios captures these uncorrelated changes between sepal and corolla (Table **2**). Descendant plants or flowers are not globally smaller, they have smaller and less conspicuous corollas. Based on the resurrection approach, patterns of trait changes between ancestral and descendant populations provide a signature of mating system and associated trait evolution. Correlation among our traits reveals that there are two clusters of highly correlated traits, floral length, labellum width and the derived floral area, thus floral shape, and rosette diameter, leaf length, leaf width and the derived leaf area, thus leaf shape (Fig. **S3**). This is why we discuss only one trait of these clusters in the following.

**Figure 1.**
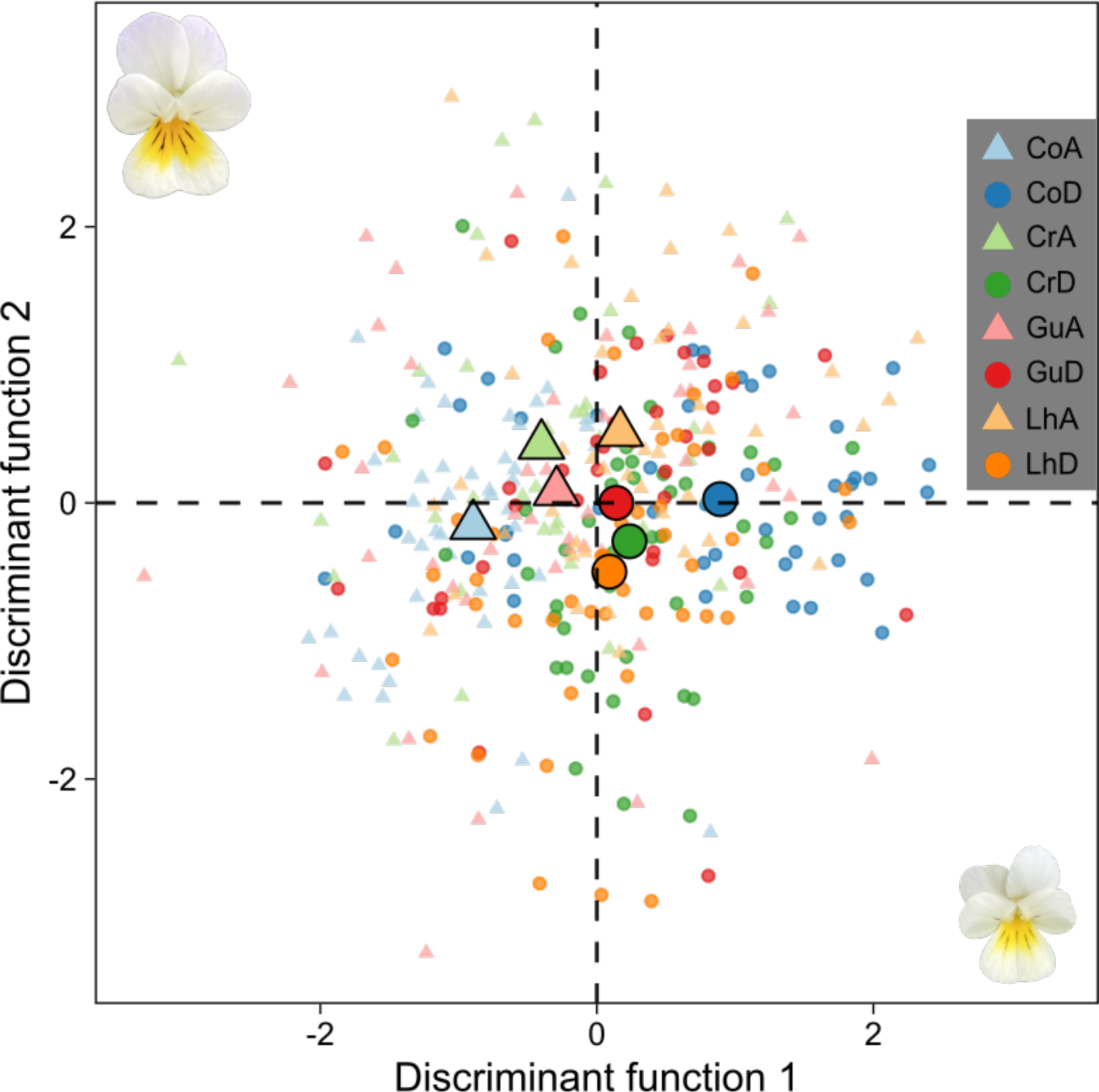
Multivariate differences between traits of ancestral and descendant populations of *Viola arvensis*. Results of a multivariate analysis (linear discriminant analysis) performed on nine traits and 357 plants (almost equally distributed among groups; N_min_ = 36 individuals; 7 points not represented because out of scale). The first two functions represent 55% and 22% of the variation respectively. The first two letters are the name of the locality (Co = Commeny; Cr = Crouy; Gu = Guernes; Lh = Lhuys). “A” (▴) ancestral population (collected in 2000 for Co, 1993 for Cr, 2001 for Gu and 1992 for Lh) and “D” (●) descendant population (all collected in 2021). Enlarged symbols are centroids of groups.

**Figure 2.**
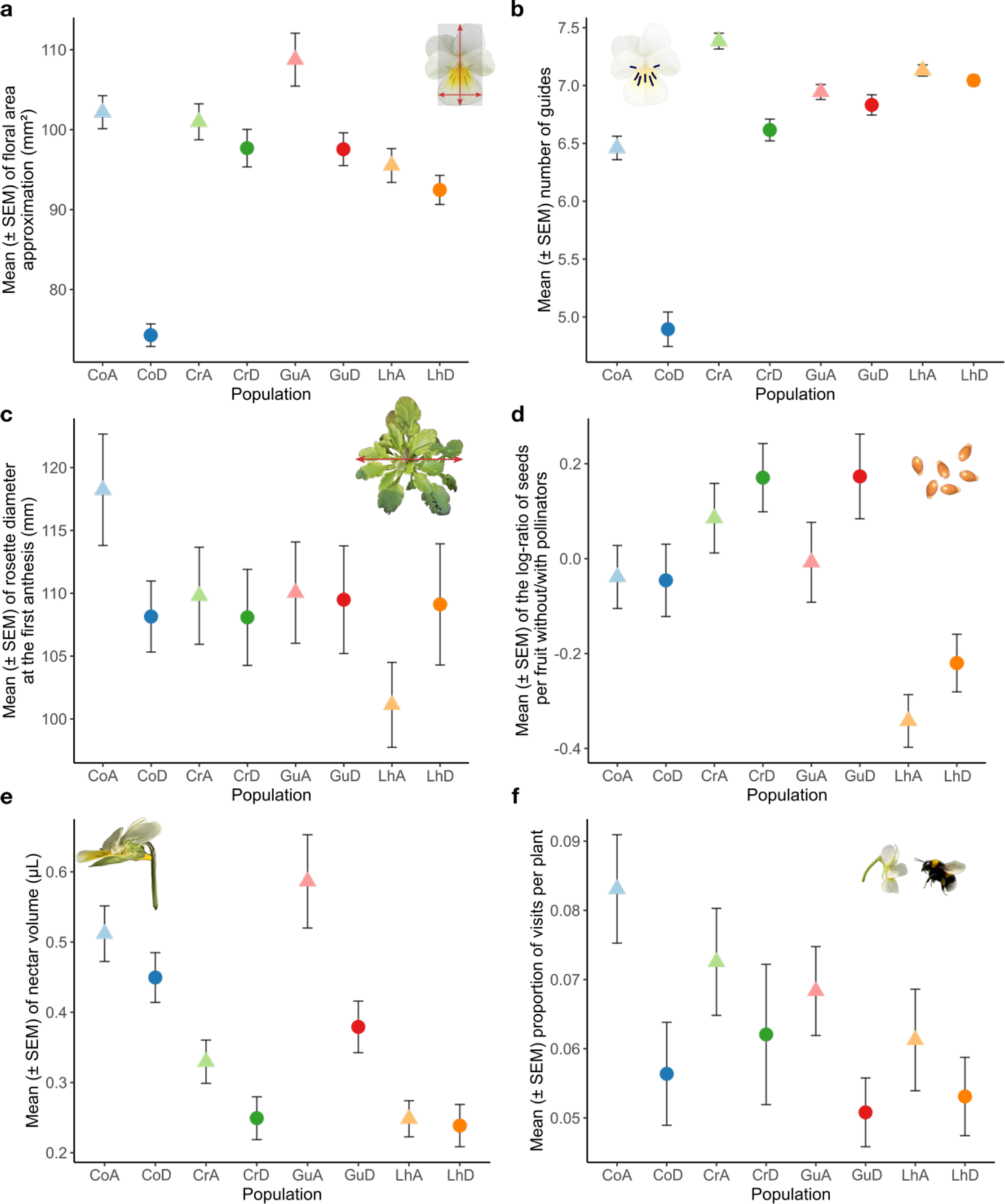
Evolutionary changes in floral, vegetative, reproductive and rewarding traits, and attractiveness of *Viola arvensis*. We measured floral traits (**a** and **b**) in the first five developed flowers per individual (N ≈ 4,000). (**a**) Floral area (multiplication of labellum width × corolla length). (**b**) Number of nectar guides. (**c**) Rosette diameter, measured on each plant at the start of flowering (N = 792). (**d**) Log-ratio of seeds produced in self-pollination compared to open pollination as a proxy of selfing ability, measured by collecting one fruit in self-pollination and one in open pollination per plant (N = 693). (**e**) Nectar production measured as the sum of the volume in three flowers per plants on fifty plants per population (N = 400). (**f**) Bumblebee preferences measured as proportion of visits per plant to a mixed plantation of 10 plants of the ancestral and 10 of the descendant populations of a single locality, exposed together to bumblebees. We recorded the number of visits to each plant by a flying bumblebee for 10 to 15 minutes in 6 to 8 replicates per location and divided it by the total number of visits during the flight (only visited plants are represented). The first two letters are the name of the locality (Co = Commeny; Cr = Crouy; Gu = Guernes; Lh = Lhuys). “A” (▴) ancestral population (collected in 2000 for Co, 1993 for Cr, 2001 for Gu and 1992 for Lh) and “D” (●) descendant population (all collected in 2021).

**Table 2.** Trait differences (mean ± SD) among our populations of *Viola arvensis*. Statistical analyses were performed using LMMs with age (ancestral or descendant), locality and the interaction between the two as fixed effects and family and, when necessary, other random effects were added (see Material and methods). When age effect was significant (bold traits), analyses were followed by two-side T-tests with a Bonferroni correction performed between the ancestral and descendant populations by locality. Significant differences in traits between ancestors and descendants (*P* < 0.05) was highlighted by bold measures and sign of changes are given using arrows (➘ for a decrease in time and ➚ for an increase in time). The first two letters are the name of the locality (Co = Commeny; Cr = Crouy; Gu = Guernes; Lh = Lhuys). “A” ancestral population (collected in 2000 for Co, 1993 for Cr, 2001 for Gu and 1992 for Lh) and “D” descendant population (all collected in 2021).

| Trait | N | CoA<br>CoD | CrA<br>CrD | GuA<br>GuD | LhA<br>LhD | Factor | Chisq | Chi<br>df | P |
| --- | --- | --- | --- | --- | --- | --- | --- | --- | --- |
| **Floral traits** |  |  |  |  |  |  |  |  |  |
| Corolla length (mm) | 3955 | **14.03 ± 1.09** | 13.28 ± 1.45 | 14.06 ± 2.18 | 13.23 ± 1.47 | Age | **15.314** | **1** | **< 0.001** |
| | | **12.09 ± 1.13** | 13.03 ± 1.88 | 13.66 ± 1.35 | 12.99 ± 1.46 | Locality | **13.384** | **3** | **0.004** |
| | | ↘ | | | | Age × Locality | **16.638** | **3** | **< 0.001** |
| Labellum width (mm) | 3955 | **7.16 ± 0.94** | 7.45 ± 0.99 | **7.51 ± 1.16** | 7.04 ± 0.76 | Age | **17.872** | **1** | **< 0.001** |
| | | **6.03 ± 0.59** | 7.30 ± 0.92 | **7.01 ± 0.88** | 6.94 ± 0.67 | Locality | **30.047** | **3** | **< 0.001** |
| | | ↘ | | ↘ | | Age × Locality | **14.725** | **3** | **0.002** |
| Floral area approximation (mm<sup>2</sup>) | 3955 | **102.2 ± 19.9** | 101.0 ± 22.5 | **108.8 ± 33.1** | 95.5 ± 21.2 | Age | **17.965** | **1** | **< 0.001** |
| | | **74.3 ± 14.0** | 97.7 ± 23.5 | **97.6 ± 20.5** | 92.5 ± 18.2 | Locality | **17.979** | **3** | **< 0.001** |
| | | ↘ | | ↘ | | Age × Locality | **14.989** | **3** | **0.002** |
| Spur length (mm) | 3954 | **4.25 ± 0.22** | 4.30 ± 0.24 | 4.30 ± 0.28 | 4.37 ± 0.19 | Age | **8.373** | **1** | **0.004** |
| | | **4.00 ± 0.23** | 4.27 ± 0.27 | 4.23 ± 0.31 | 4.38 ± 0.23 | Locality | **42.226** | **3** | **< 0.001** |
| | | ↘ | | | | Age × Locality | **12.388** | **3** | **0.006** |
| Sepal length (mm) | 3955 | 13.92 ± 1.32 | 14.26 ± 1.51 | 12.64 ± 1.34 | 13.61 ± 1.19 | Age | 0.550 | 1 | 0.459 |
| | | 13.55 ± 1.01 | 13.95 ± 1.51 | 12.71 ± 1.35 | 13.77 ± 1.32 | Locality | **63.509** | **3** | **< 0.001** |
| | | | | | | Age × Locality | 2.7607 | 3 | 0.430 |
| Sepal/corolla length ratio | 3955 | **0.998 ± 0.078** | 1.088 ± 0.117 | 0.918 ± 0.120 | **1.044 ± 0.102** | Age | **14.137** | **1** | **< 0.001** |
| | | **1.132 ± 0.099** | 1.093 ± 0.125 | 0.941 ± 0.107 | **1.077 ± 0.083** | Locality | **93.828** | **3** | **< 0.001** |
| | | ↗ | | | ↗ | Age × Locality | **15.302** | **3** | **0.002** |
| Number of guides | 3993 | $6.46 \pm 0.98$ | $7.38 \pm 0.68$ | $6.94 \pm 0.64$ | $7.13 \pm 0.48$ | Age | 33.591 | 1 | < 0.001 |
| | | $4.89 \pm 1.48$ | $6.62 \pm 0.94$ | $6.83 \pm 0.88$ | $7.04 \pm 0.42$ | Locality | 112.889 | 3 | < 0.001 |
|  |  | ↘ | ↘ |  |  | Age × Locality | 30.675 | 3 | < 0.001 |
| Anthesis duration (days) | 3991 | $6.39 \pm 0.85$ | $6.20 \pm 1.06$ | $7.85 \pm 1.56$ | $7.00 \pm 1.04$ | Age | 9.261 | 1 | 0.002 |
| | | $5.07 \pm 0.85$ | $6.07 \pm 0.91$ | $7.27 \pm 1.29$ | $7.41 \pm 0.81$ | Locality | 137.555 | 3 | < 0.001 |
|  |  | ↘ |  | ↘ | ↗ | Age × Locality | 24.407 | 3 | < 0.001 |
| Flowering date (days after the first flowering) | 792 | $20.31 \pm 6.78$ | $17.48 \pm 8.78$ | $19.58 \pm 10.81$ | $19.30 \pm 7.57$ | Age | 0.027 | 1 | 0.870 |
| | | $18.54 \pm 4.58$ | $17.57 \pm 7.82$ | $20.23 \pm 12.69$ | $19.69 \pm 9.13$ | Locality | 3.783 | 3 | 0.286 |
|  |  |  |  |  |  | Age × Locality | 1.007 | 3 | 0.800 |
| Average floral display (open flowers per day) | 789 | $6.37 \pm 2.39$ | $6.85 \pm 2.59$ | $7.89 \pm 3.47$ | $6.64 \pm 2.31$ | Age | 8.190 | 1 | 0.004 |
| | | $4.64 \pm 1.99$ | $6.89 \pm 2.87$ | $6.95 \pm 3.26$ | $6.43 \pm 2.17$ | Locality | 32.182 | 3 | < 0.001 |
|  |  | ↘ |  |  |  | Age × Locality | 7.855 | 3 | 0.049 |
| Vegetative traits |  |  |  |  |  |  |  |  |  |
| Rosette diameter (mm) | 792 | $118.23 \pm 42.75$ | $109.80 \pm 38.60$ | $110.05 \pm 40.29$ | $101.12 \pm 33.77$ | Age | 0.194 | 1 | 0.660 |
| | | $108.15 \pm 28.11$ | $108.08 \pm 38.24$ | $109.49 \pm 42.85$ | $109.11 \pm 48.21$ | Locality | 5.601 | 3 | 0.133 |
|  |  |  |  |  |  | Age × Locality | 2.397 | 3 | 0.494 |
| Leaf length (mm) | 792 | $26.30 \pm 6.38$ | $24.10 \pm 5.63$ | $22.34 \pm 5.18$ | $21.57 \pm 4.85$ | Age | 1.226 | 1 | 0.268 |
| | | $22.42 \pm 3.95$ | $24.28 \pm 5.65$ | $23.17 \pm 6.07$ | $21.94 \pm 5.63$ | Locality | 19.895 | 3 | < 0.001 |
|  |  |  |  |  |  | Age × Locality | 11.311 | 3 | 0.010 |
| Leaf width (mm) | 792 | $19.84 \pm 3.77$ | $18.67 \pm 3.03$ | $17.02 \pm 2.76$ | $17.58 \pm 3.07$ | Age | 2.499 | 1 | 0.114 |
| | | $17.96 \pm 2.56$ | $18.94 \pm 3.01$ | $16.66 \pm 2.91$ | $17.66 \pm 3.07$ | Locality | 40.014 | 3 | < 0.001 |
|  |  |  |  |  |  | Age × Locality | 8.024 | 3 | 0.046 |
| Leaf area approximation (mm²) | 792 | $541.3 \pm 231.9$ | $461.3 \pm 173.6$ | $391.3 \pm 152.8$ | $390.2 \pm 147.1$ | Age | 1.910 | 1 | 0.167 |
| | | $410.5 \pm 128.7$ | $472.4 \pm 177.6$ | $398.5 \pm 173.7$ | $401.6 \pm 173.0$ | Locality | 27.414 | 3 | < 0.001 |
|  |  |  |  |  |  | Age × Locality | 10.559 | 3 | 0.014 |
| Biomass (g) | 777 | $7.243 \pm 2.515$ | $8.220 \pm 2.712$ | $8.027 \pm 2.624$ | $8.090 \pm 2.901$ | Age | 0.029 | 1 | 0.864 |
| | | $8.034 \pm 2.553$ | $7.789 \pm 2.747$ | $7.606 \pm 2.904$ | $8.386 \pm 3.066$ | Locality | 3.490 | 3 | 0.322 |
|  |  |  |  |  |  | Age × Locality | 4.850 | 3 | 0.183 |
| Reproductive traits |  |  |  |  |  |  |  |  |  |
| Log-ratio of the number of seeds without/with pollinators | 696 | $-0.039 \pm 0.619$ | $0.085 \pm 0.659$ | $-0.008 \pm 0.775$ | $-0.342 \pm 0.546$ | Age | 3.257 | 1 | 0.071 |
| | | $-0.046 \pm 0.711$ | $0.171 \pm 0.678$ | $0.173 \pm 0.757$ | $-0.220 \pm 0.601$ | Locality | 38.808 | 3 | < 0.001 |
|  |  |  |  |  |  | Age × Locality | 1.674 | 3 | 0.643 |
| Weight per seed with pollinators (mg) | 659 | $0.446 \pm 0.199$ | $0.421 \pm 0.153$ | $0.301 \pm 0.154$ | $0.541 \pm 0.146$ | Age | 4.984 | 1 | 0.026 |
| | | $0.385 \pm 0.161$ | $0.367 \pm 0.145$ | $0.300 \pm 0.143$ | $0.546 \pm 0.142$ | Locality | 213.732 | 3 | < 0.001 |
|  |  |  |  |  |  | Age × Locality | 6.197 | 3 | 0.102 |
| Weight per seed without<br>pollinators (mg) | 704 | **0.689 ± 0.206** | 0.633 ± 0.169 | 0.517 ± 0.187 | 0.764 ± 0.137 | Age | **11.884** | **1** | **< 0.001** |
|  |  | **0.565 ± 0.200** | 0.584 ± 0.162 | 0.501 ± 0.189 | 0.788 ± 0.117 | Locality | **240.861** | **3** | **< 0.001** |
|  |  | ↘ |  |  |  | Age × Locality | **22.227** | **3** | **< 0.001** |
| ***Plant-pollinator interaction traits*** |  |  |  |  |  |  |  |  |  |
| Nectar volume (μL) | 400 | 0.512 ± 0.280 | **0.330 ± 0.218** | **0.586 ± 0.470** | 0.248 ± 0.182 | Age | **12.605** | **1** | **< 0.001** |
|  |  | 0.450 ± 0.251 | **0.249 ± 0.215** | **0.379 ± 0.260** | 0.239 ± 0.213 | Locality | **75.120** | **3** | **< 0.001** |
|  |  |  | ↘ | ↘ |  | Age × Locality | 4.636 | 3 | 0.200 |
| Bumblebee visit preferences<br>(proportion of visit per plant) | 456 | **0.083 ± 0.008** | 0.073 ± 0.008 | 0.068 ± 0.006 | 0.061 ± 0.007 | Age | **14.722** | **1** | **< 0.001** |
|  |  | **0.056 ± 0.007** | 0.062 ± 0.010 | 0.051 ± 0.005 | 0.053 ± 0.006 | Locality | 2.027 | 3 | 0.567 |
|  |  | ↘ |  |  |  | Age × Locality | 1.950 | 3 | 0.583 |

Regarding reproductive traits, our results revealed a marginally significant (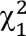 = 3.257, *P* = 0.071) increase over time in the ability to set seeds in the absence of pollinators (Fig. **2d**). The log-ratio can be greater than zero because fruits higher in the stem, open pollinated fruits in our experimental design, produce less seeds, independently to pollinators. However, we found a significant decrease in seed weight in open- and self-pollination (Table **2**).

Our study also reveals that both locality and the interaction between locality and age (ancestors or descendants) have a strong effect on trait variations. The locality effects indicate spatial differentiation in measured traits, whereas the locality×age interaction reveals a site dependent evolution. The latter interaction effects were different in magnitude, but not in sign, between ancestors and descendants for all the traits, except anthesis duration (Table **2**). Hence, evolution in the four localities exhibits the same directional change *i.e.* evidence of convergent shifts towards a selfing syndrome in the four independent populations (Fig. **1**) while differing in magnitude.

We found a consistent decrease of 20% on average in rewarding traits, as represented by measurements of nectar volume (Fig. **2e**, Table **2**).

We assessed how this evolution impacts plant-pollinator interactions from the point of view of pollinators through a preference experiment involving visits of the bumblebees *Bombus terrestris* to a mixture of ancestral and descendant plants. Bumblebees visited ancestral plants more frequently than they visited descendant plants, showing a preference towards the former that is consistent with their higher attractiveness and reward traits (Fig. **2f**, Table **2**).

## Discussion

Overall, we found decreases in plant-insect mediating traits such as floral display, floral area, nectar guides, which are assumed to promote recognition of the path to floral rewards (Hansen *et al*., 2012), and nectar production. The lack of consistent changes in vegetative traits, except for leaf traits in Commeny which could be explained by the earlier flowering (Fig. **S3**), leads us to conclude that changes observed are not due to inbreeding depression, especially because late acting traits such as biomass exhibit no changes (Winn *et al*., 2011). Moreover, in an additional experiment, we have measured a very low inbreeding depression on floral traits (around 3%). Such a value, far below the observed trait shift, cannot account for the observed shift (*in prep*). Changes in sepal to corolla length ratio but not in sepal length (Table **2**) point out evidence of differential evolution of petal and sepal lengths despite their homologous origins (Breuninger & Lenhard, 2010). This provides strong evidence for selection against specific alleles coding costly traits that mediate interactions with pollinators.

Decreases in plant-insect mediating traits (e.g., floral area, nectar production) are well accompanied by decreases in floral attractiveness, as well as an increase in selfing rates. This is also accompanied by an evidence for enhanced selfing ability, except for Commeny and Crouy in agreement with the lack of significantly increased selfing rate. Decrease in seed weight, characterizing maternal investment in individual progeny, is consistent with a shift towards increased reproductive assurance associated with reduced seed investment, as observed in previous studies (De Jong *et al*., 2005). These changes are all consistent with the ongoing evolution of a selfing syndrome (Ornduff, 1969; Sicard & Lenhard, 2011). Furthermore, we found such patterns to be convergent across the four populations investigated (Fig. **1**), despite changes in magnitude. Such changes in magnitude can be due to genetic drift, differences in genetic variances for the traits or differences in selective pressures among localities.

The absence of significant change in selfing ability and selfing rate in Commeny despite the highest change in floral traits could be explained by an earlier evolution toward selfing than in other populations. Indeed, we only capture a small part of the evolution trajectory toward selfing syndrome. We may suppose that traits directly linked to selfing (herkogamy or dichogamy for example) evolved earlier than traits mediating plant-pollinator interaction, linked to investment in outcrossing (this pattern is also observed in Bodbyl Roels & Kelly, 2011). Following this hypothesis, evolution of traits mediating plant-pollinator interaction is secondary and driven by the release of the selective pressure imposed by pollinators, the cost of maintenance of traits and potential reallocations. In this case, selfing ability or selfing rate could have already evolved in ancestral population (pollinator decline has been recorded since 1970, Biesmeijer *et al*., 2006), resulting in slight changes between our two samples (Fig. **2d**, Table **1**). With increased selfing, floral traits linked to investment in outcrossing are just costs which can experience large changes (Fig. **2a-b**). Earlier evolution is consistent with the fact that the ancestral population of Commeny had the highest selfing rate of ancestral populations (Table **1**).

In spite of these local differences, this study thus provides a unique demonstration for a convergent evolutionary change in plant mating systems over a short-time scale in natural populations. Indeed, despite similar floral trait evolutions were found in a single population of *Volia arvensis* in another French intensive agricultural area (Cheptou *et al*., 2022), this study is the first to show convergent evolution across populations, reduced rewarding trait, and reduced attractiveness.

The decrease in nectar production in our study species provides evidence of a reduced interaction between plants and their pollinators. Indeed, floral nectar production is costly (Pyke, 1991) and only dedicated to rewarding pollinators, thus is key to the maintenance of stable plant-pollinator interactions (Brandenburg *et al*., 2012). While angiosperm-pollinator interactions have evolved over the long-term, our study shows that current environmental changes can drive a rapid evolution towards a breakdown of such interactions.

Recurrent evolution towards selfing has been repeatedly documented in the wild, but to our knowledge, only at the phylogenetic time scale (Stebbins, 1957; Barrett, 2002). This evolutionary transition is classically considered to be irreversible (the “selfing as an evolutionary dead-end” hypothesis (Stebbins, 1957; Igic & Busch, 2013)) and higher extinction rates of such linages have been reported (Goldberg *et al*., 2010). Evolution towards selfing could thus be driven by natural selection over the short-term but could impede long-term plant population survival (Cheptou, 2019). The increase in selfing rate is indeed expected to have genomic impacts on individuals and populations through inbreeding (homozygosity), reduced effective population sizes and increased genetic drift (Busch *et al*., 2022). Although our results do not support this, it is interesting to note that the only location showing reduced genetic richness, Commeny, is also the one showing the most pronounced change in morphology and thus experienced the more pronounced selection. This reduction of genetic richness could be the product of drift and of the interaction of inbreeding and selection (genetic draft, Busch *et al*., 2022). The absence of reduction in genetic diversity in other locations could be due to large enough population size and thus very limited effect of genetic drift or recent increase in inbreeding consistent with low morphology differentiation. However, linked to a decreased genetic diversity, selfer populations may have a reduced potential response to future selective pressures (Glémin & Ronfort, 2013; Noël *et al*., 2017). The short-term evolution of selfing in *Viola arvensis* we document here is no-doubt facilitated by its initial mixed mating system (contrary to self-incompatible plants for example, Thomann *et al*., 2015). Nonetheless our results illustrate how mixed-mating species may evolve towards a higher reliance on self-pollination and reduced plant-pollinator interactions in the face of global change, in particular pollinator declines. If this rapid transition towards a selfing syndrome reflects a broader trend among Angiosperms, it may reflect a concerning extinction debt (Goldberg *et al*., 2010).

Ongoing environmental changes – such as habitat destruction and fragmentation, agricultural land pollution and species’ introductions – have been documented as causes of pollinator declines, both by affecting pollinator populations directly (Potts *et al*., 2010; Goulson *et al*., 2015), and indirectly through their impacts on plant community composition, affecting the quantity and diversity of nectar resources (Goulson *et al*., 2015; Baude *et al*., 2016). In addition, and as we illustrate here, floral nectar production can also evolve rapidly in plant species in response to a relaxed selective pressure associated with pollinator declines, prompting declines in nectar resources of a magnitude similar to those associated with changes in plant community composition (Baude *et al*., 2016). These decreases in nectar production may then reinforce pollinator declines if nectar levels fall below those necessary to sustain wild bee populations. Environmental changes may thus present a double jeopardy to pollinator populations, as they become victims of both the changes themselves and of plant trait evolution (Weinbach *et al*., 2022). This in turn may result in an eco-evolutionary positive feedback loop that furthers pollinator declines, further reinforcing plant evolution towards a selfing syndrome. This can explain plant-pollinator network degradation as documented in a previous study (Burkle *et al*., 2013) and raises the concerning prospect of cascading effects in trophic networks in general, beyond plant-pollinator interactions.

In summary, our study highlights the potential of natural populations to respond quickly to environmental changes. However, such evolutionary responses may have impacts on ecological interactions, here plant-pollinator interactions, and potentially cascading trophic consequences in ecosystems. There is thus an urgent need to investigate whether these results are symptomatic of a broader pattern among angiosperms and their pollinators, and if so understanding whether there is a possibility to reverse this process and break this eco-evolutionary positive feedback loop.

## Acknowledgments

We thank David Delguerde for construction and installation of insect-proof cages, Pauline Durbin and Fabien Lopez for help in taking care of the plants during the experiment, Marie-Pierre Dubois for lab-work guidance, the Conservatoire Botanique National de Bailleul and the Conservatoire Botanique National du Bassin Parisien for the supply of seeds and information on sampling locations, Anaïs Vignerol for help in data collection, Ana Rodrigues, John D. Thompson, Pierre-André Crochet and three anonymous reviewers for improving this manuscript. Genotyping data was obtained using technical facilities of GenSeq of the “Institut des Sciences de l’Evolution de Montpellier” with the support of LabEx CeMEB an ANR “Investissements d’avenir” program (ANR-10-LABX-04-01).

## Author contributions

Conceptualization: POC, SAP; methodology: SAP, POC; lab work: LD, SAP; data collection: SAP, PG, ADM, VP, POC; analysis: SAP, LD; writing – original draft: SAP; writing – review & editing: SAP, POC

## Data Availability Statement

The data that support the findings of this study are available in data.InDoRES at https://doi.org/10.48579/PRO/VIJEHR.

## Supporting information

**Fig. S1** Graphical experimental setup

**Fig. S2** Description of morphometric measurements

**Fig. S3** Correlation between measured traits

**Table S1** Information on sampled populations

