## Supplementary information for "Ongoing convergent evolution of a selfing syndrome threatens plant-pollinator interactions"

**Supporting information**

**Fig. S1** Graphical experimental setup

**Fig. S2** Description of morphometric measurements

**Fig. S3** Correlation between measured traits

**Table S1** Information on sampled populations

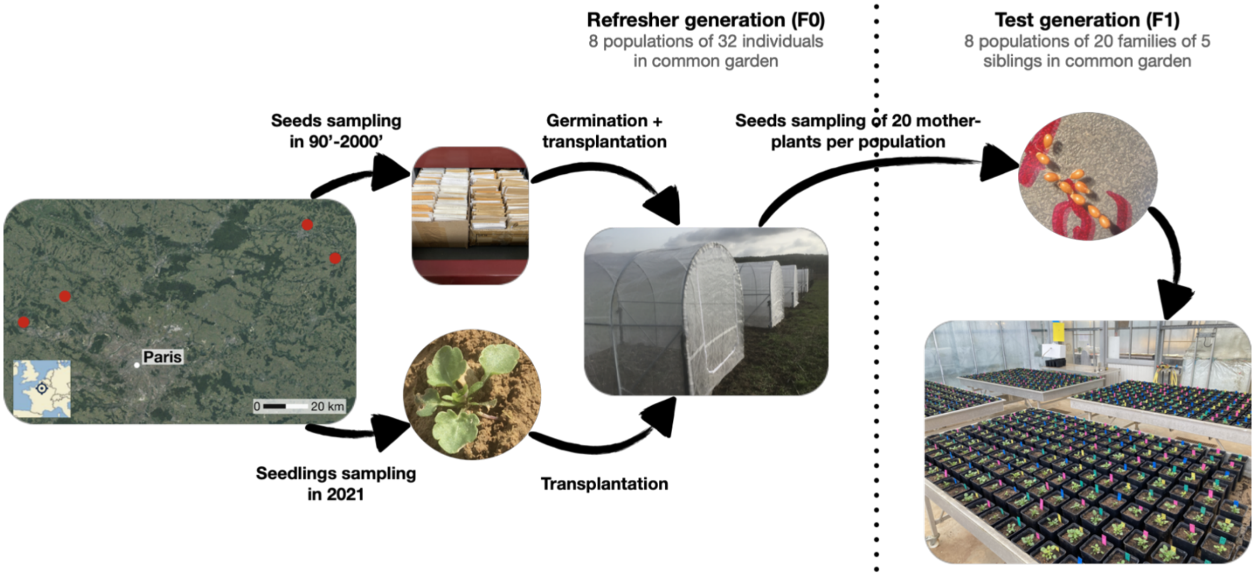


**Fig. S1.** Experimental design. Four localities were sampled two times, with a minimum 20 years between the two samplings. The refresher generation (F0) was performed in common garden. It was used to analyse population genetics and to produce a family design of 20 families of 5 siblings per population for the test generation (F1), on which we measured traits.

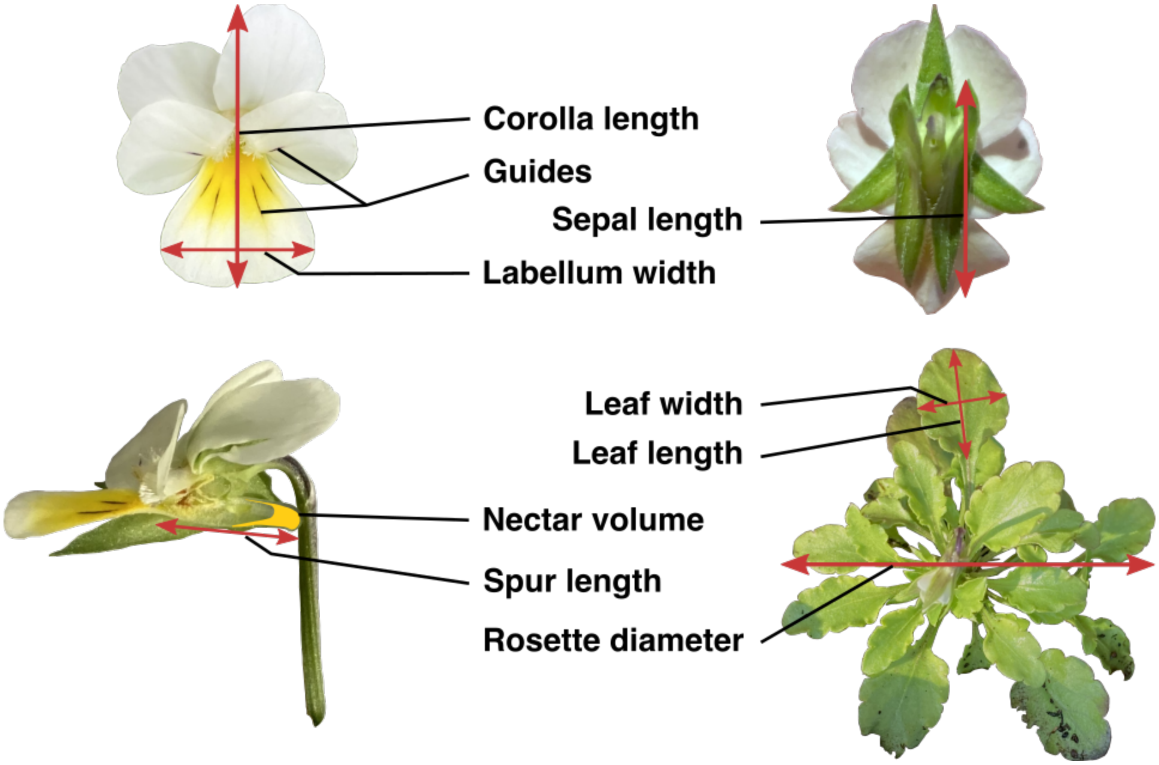


**Fig. S2.** Trait measures. List of traits measured and delimitation of their measurements. Traits represented in this figure are only traits which were directly measured on plants.

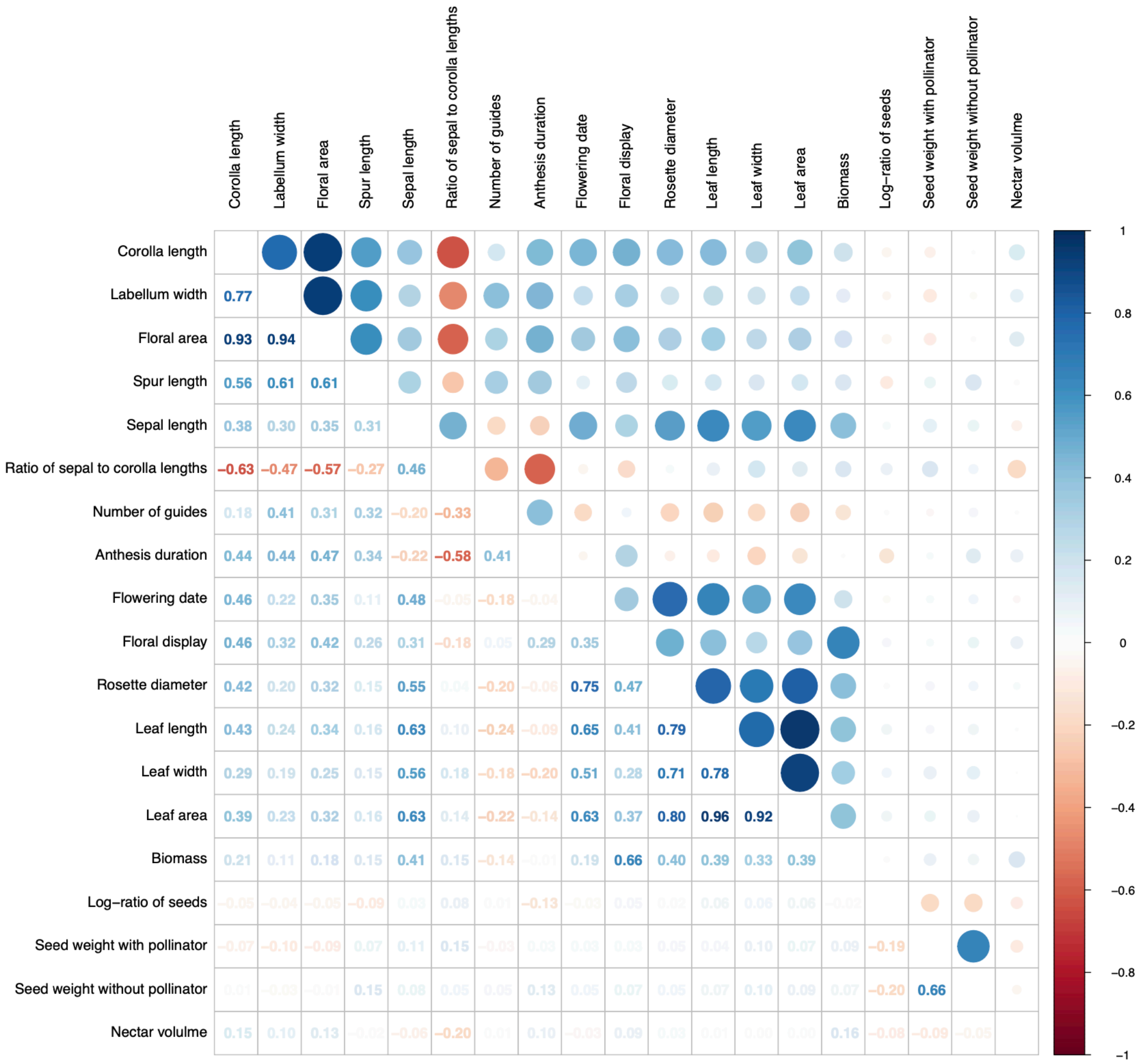


**Fig. S3.** Correlation between measured traits presented in Table **2**.

**Table S1.** Position and year of the sampling of our populations.

| *Location* | **Commeny** (49°7'37.11"N, 1°53'46.30"E) | **Crouy** (49°24'24.31"N, 3°22'20.73"E) | **Guernes** (49° 1'10.75", 1°38'46.82"E) | **Lhuys** (49°16’35.40"N, 3°32’58.52"E) |
|---|---|---|---|---|
| **Year of first sampling (ancestors)** | 2001 | 1993 | 2000 | 1992 |
| **Year of second sampling (descendants)** | 2021 | 2021 | 2021 | 2021 |
